# Pectin methylesterification modulates cell wall properties to promote neighbour proximity-induced hypocotyl growth

**DOI:** 10.1101/2023.03.07.531598

**Authors:** Fabien Sénéchal, Sarah Robinson, Evert Van Schaik, Martine Trévisan, Prashant Saxena, Didier Reinhardt, Christian Fankhauser

**Affiliations:** Centre for Integrative Genomics, Faculty of Biology and Medicine, Génopode Building, University of Lausanne, Lausanne, Switzerland; Institute of Plant Sciences, University of Bern, Bern, Switzerland; Department of Biology, University of Fribourg, Fribourg, Switzerland

## Abstract

Plants growing with neighbours compete for light and consequently increase growth of their vegetative organs to enhance access to sunlight. This response, called shade avoidance syndrome (SAS), involves photoreceptors such as phytochromes as well as phytochrome interacting factors (PIFs), which regulate the expression of growth-mediating genes. Numerous cell wall-related genes belong to the putative targets of PIFs, and the importance of cell wall modifications for enabling growth was extensively shown in developmental models such as dark-grown hypocotyl. However, the role of the cell wall in the growth of de-etiolated seedlings regulated by shade cues remains poorly established. Through analyses of mechanical and biochemical properties of the cell wall coupled with transcriptomic analysis of cell wall-related genes, we show the importance of cell wall modifications in neighbour proximity-induced elongation. Further analysis using loss-of-function mutants impaired in the synthesis and remodeling of the main cell wall polymers corroborated this. We focused on the *cgr2cgr3* double mutant that is defective in homogalacturonan (HG) methyltransferase activity required for methylesterification of HG-type pectins. By following hypocotyl growth kinetically and spatially and analyzing the mechanical and biochemical properties of cell walls, we found that methylesterification of HG-type pectins was required to enable global cell wall modifications. Moreover, HG-class pectin modification was needed for plant competition-induced hypocotyl growth. Collectively our work suggests that in the hypocotyl PIFs orchestrate changes in the expression of numerous cell wall genes to enable neighbour proximity-induced growth.

**One sentence summary:** The degree of methylesterification of pectins modulates global changes in the cell wall and its mechanical properties that contribute to the neighbour proximity-induced hypocotyl growth in Arabidopsis

## Introduction

Plants growing in dense populations compete for light required for photosynthesis (Fiorucci and Fankhauser, 2017). Proximity of competitors and shade are perceived as a change in light intensity and quality, resulting in various developmental adaptations (Galvão and Fankhauser, 2015). Depending on their response to shade, plants are classified into shade-tolerant and shade-avoiding species (Gommers et al., 2013). The latter react to shade with characteristic growth responses in order to reach full sunlight for photosynthesis. This phenomenon, known as Shade-Avoidance Syndrome (SAS) can be observed in most aerial organs and involves a range of developmental changes (Pierik and De Wit, 2014). For example, SAS entails early flowering and inhibition of branching, leaves adopt an upright position (hyponasty), and petioles, stems and hypocotyls elongate (de Wit et al., 2016). These changes accelerate the life cycle of plants and warrant survival and propagation under limited light availability.

The phytochrome (phy) type photoreceptors have a central role in SAS with phyB playing a predominant role in *Arabidopsis thaliana* (Legris et al., 2019). Under high R:FR conditions, corresponding to full sunlight, active phyB moves into the nucleus, where it interacts with transcription factors known as Phytochrome Interacting Factors (PIFs) to inhibit their activities. Low R:FR conditions, on the other hand, cause inactivation of phyB and derepression of the PIFs, which modulate genes required for shade-induced growth (de Wit et al., 2016). A central mechanism in shade-induced growth is auxin biosynthesis in cotyledons and young leaves followed by polar transport and distribution in hypocotyls and stems, which elongate in response to shade (de Wit et al., 2014). phyB and PIFs also act localy in the hypocotyl to promote growth through mechanisms that are less clearly established (Fiorucci and Fankhauser, 2017; Pucciariello et al., 2018). For example PIFs regulate the expression of genes required for plasma-membrane lipid biogenesis in the hypocotyl (Ince et al., 2022). Moreover, numerous genes encoding cell wall-modifying proteins are induced by shade and targeted by PIFs (Kohnen et al., 2016; Pedmale et al., 2016), suggesting role for cell wall metabolism in the establishment of the shade-regulated growth.

The primary cell wall of growing plant organs has seemingly contradictory functions. On the one hand, it provides mechanical strength to maintain cell shape and plant stature, on the other hand, it has to remain elastic and plastic to allow cell expansion and plant growth (Bashline et al., 2014). In dicotyledonous species such as Arabidopsis, the primary cell wall consists of interconnected networks of polysaccharides and structural proteins/glycoproteins (Cosgrove, 2005; Wolf et al., 2012a; Nguema-Ona et al., 2014). A first network of cellulose microfibrils cross-linked by hemicelluloses (network 1) is embedded in a second network made by pectins that have gelling properties (network 2). Pectins are intimately associated with a third network (network 3), consisting mainly of glycoproteins such as extensins (EXTs) and arabinogalactan proteins (AGPs) (Hijazi et al., 2014). This complex three-dimensional mesh resists to internal turgor pressure, and at the same time yields to allow cell growth. Dynamic adjustment of physical cell wall properties (e.g. stiffness, elasticity, plasticity) involves cell wall-synthesizing and modifying enzymes that usually belong to multigenic families and constantly modulate cell walls to allow growth (Atmodjo et al., 2013; Sénéchal et al., 2014b; Pauly and Keegstra, 2016; Showalter and Basu, 2016).

Cell wall remodeling plays a central role in the control of seedling development, however, its contribution to adaptive growth phenomena in response to environmental cues such as SAS remains poorly understood. Although transcriptomic analyses suggested that cell wall remodeling may play a central role in adaptive growth processes (Kohnen et al., 2016), direct functional evidence is scarce. A role for xyloglucans in SAS has been established for petiole elongation in Arabidopsis (Sasidharan et al., 2010; Sasidharan and Pierik, 2010), and for shade-induced growth in *Stellaria longipes* (Sasidharan et al., 2008). Here, we took a systematic approach to explore the contribution of cell wall remodeling in SAS of the Arabidopsis hypocotyl. By combining Fourier-transformed infrared (FTIR) analyses of cell wall constituents with measurements of cell wall biophysics, we show that cell wall remodeling is triggered at early stages of the growth response induced by low R:FR, indicative of neighbour proximity that is a form of SAS. Employing systematic transcriptomic and genetic analysis with mutants affected in various aspects of cell wall biosynthesis and modification, we establish pectin methylesterification status as a central determinant of cell wall extensibility in the neighbour proximity-induced hypocotyl growth.

## Results

### Low R:FR induces cell elongation mainly in the middle part of the hypocotyl

In order to simulate the proximity of competing neighbours, we subjected Arabidopsis seedlings to white light supplemented with far red (low R:FR ratio). Within one day, this led to increased hypocotyl growth compared to control seedlings kept in white light (high R:FR ratio) (**Fig. 1A**). In order to determine the site of growth, we used the borders of epidermal cells as marks, since hypocotyl elongation proceeds almost exclusively by cell elongation without cell division. In order te assess cell dimensions, confocal images of seedlings were segmented in MorphoGraphX (de Reuille et al., 2015), followed by semiautomated cell size measurements. Cell length along the length of the hypocotyl was increased by low R:FR primarily in the middle part of the hypocotyl (**Fig. 1B**).

**Figure 1.**
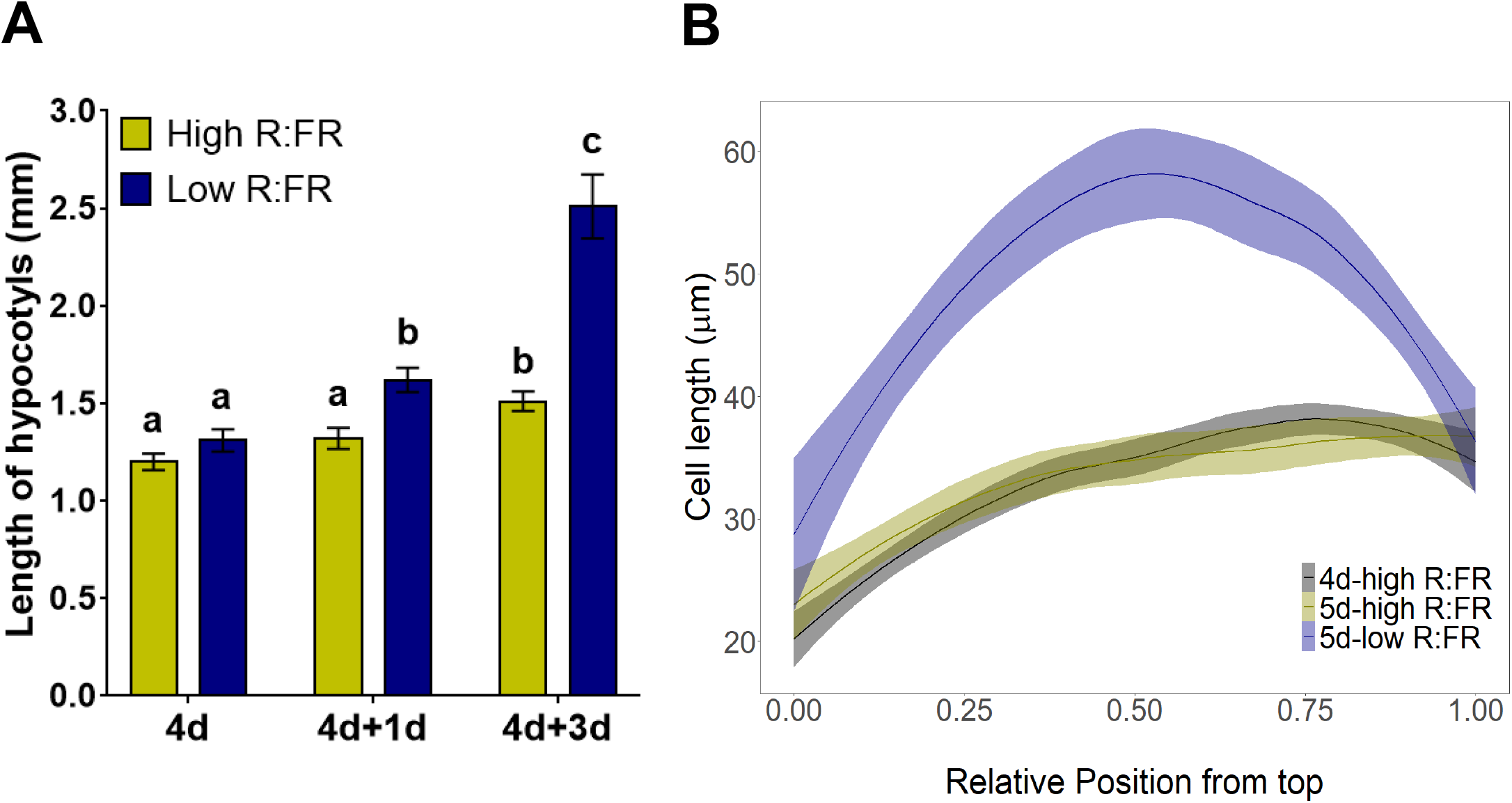
Low R:FR induces fast growth in the middle part of the hypocotyl. **(A)** Length of the hypocotyls in response to high and low R:FR treatments. Col-0 seedlings were grown for 4 days in high R:FR before being transferred under low R:FR (blue bars) or kept in high R:FR (yellow bars) for 3 additional days. The bars show means in mm ± confidence intervals measured at 4 (4d), 5 (4d+1d) and 7 (4d+3d) days. Significant differences (indicated with letters) were determined according to one-way Anova followed by a multiple comparisons with Tukey’s test. **(B)** Length of the hypocotyl epidermis cells in response to high and low R:FR treatments. After 4 days under high R:FR (blue curve), Col-0 seedlings were grown for 1 day under high R:FR (red curve) or low R:FR (green curve). The length of the epidermis cells were plotted from the top to the bottom part of the hypocotyl. The curves show means in µm ± confidence intervals (shaded areas).

### Low R:FR changes mechanical properties of hypocotyl cell walls

To assess the mechanical properties of the hypocotyl during growth induced by low R:FR, we used an automated confocal micro-extensometer (Robinson et al., 2017). This approach enabled *in vivo* quantification of the elastic properties of cell walls. Hypocotyls of intact seedlings were abraded by freezing and thawing and subjected to repeated cycles of application and removal of 5 mN of force. Our analysis revealed increased strain under low R:FR after one day of treatment, and decreased strain after three days (**Fig. 2A**). Interestingly this effect was specific to low R:FR conditions, since no such changes were observed under high R:FR (**Fig. 2A**).

**Figure 2.**
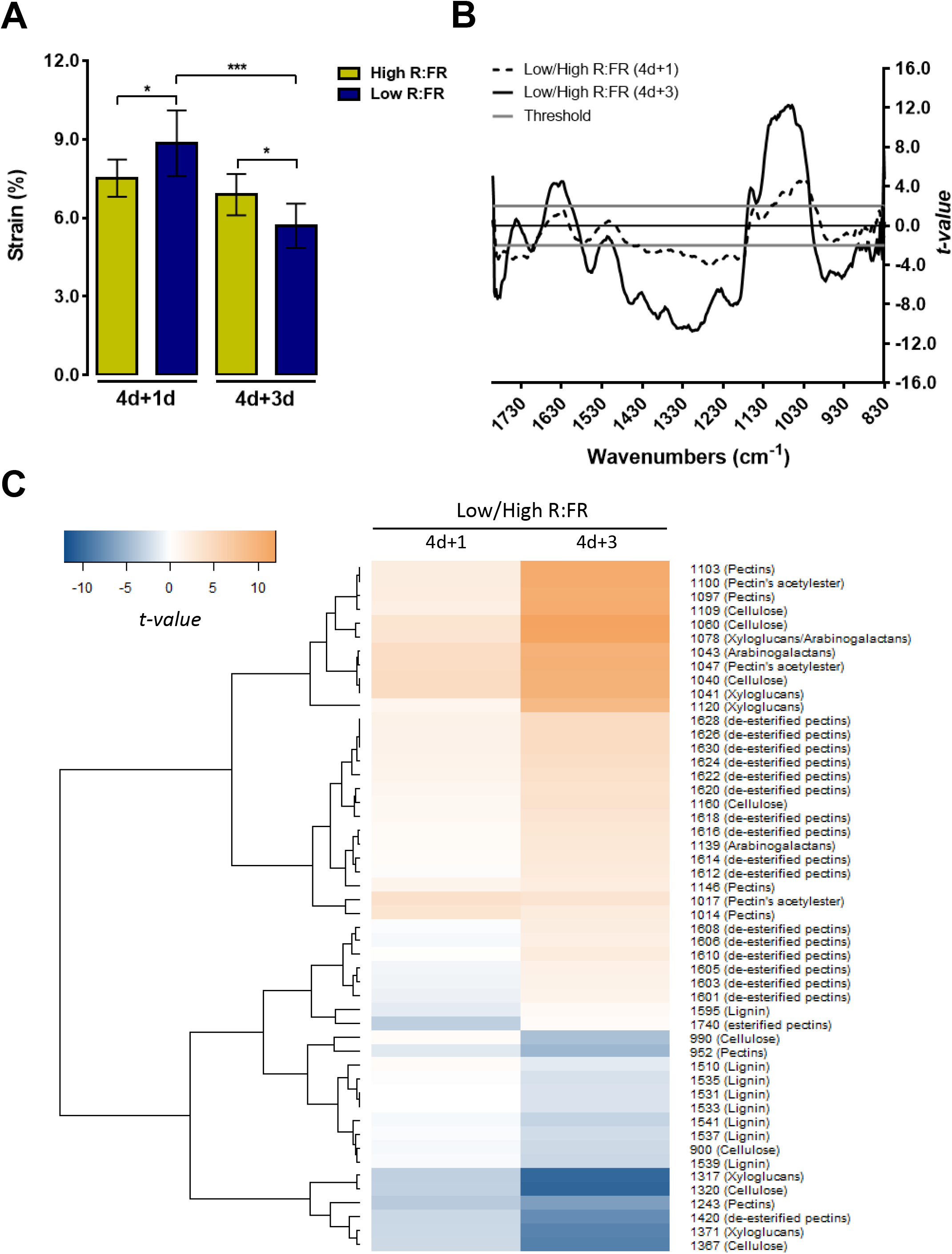
Low R:FR induces changes in mechanical and cell wall properties of the hypocotyl. **(A)** Elastic properties of hypocotyl assessed under high and low R:FR treatment. Hypocotyls were frozen and thawed then subjected to cycles of application and removal of 5 mN of force using an automated confocal micro extensometer (see methods). The average magnitude of strain incurred by seedlings grown in a high R:FR (yellow bars) or low R:FR (blue bars) light regime after 1 (4d+1d) and 3 days (4d+3d) is shown. Bright-field images were collected every 645 ms and strain was computed from regions that were tracked in the images using the ACME tracker software. The bars show means in % ± SD (n>10 independent seedlings, at least five oscillations were made). Pairwise comparisons were made using Welch t-test brackets indicated statistical tests that were made with significance p<0.1*, p<0.05** and p<0.01***. **(B**,**C)** Cell wall properties in the middle part of the hypocotyl under high and low R:FR treatments. Cell wall chemical bounds were analyzed by Fourrier-Transformed InfraRed (FTIR) microspectroscopy. For each hypocotyl, 6 spectra were collected in the middle part, avoiding the central cylinder, for at least 5 independent hypocotyls per condition. Baseline correction and data normalization were made for the absorbances between 1810 and 830 cm^-1^ (corresponding to the cell wall fingerprint, **see Supplemental Figure S1**). Pairwise comparison between high and low R:FR was made after 1 and 3 days treatments and significant differences were identified using Student’s t-test for each wavelength. **(B)** All Student’s t-values were plotted against wavelengths with horizontal lines referring to significant threshold for p<0.05. Student’s t-values above +2 or below -2 indicate respectively an enrichment or an impoverishment of cell wall components in low compared to high R:FR. **(C)** Student’s t-values for wavelengths assigned to cell wall components were used to build the heatmap with negative and positive t-values respectively represented by a range of colors from blue to orange.

To relate growth and mechanical changes induced by low R:FR to cell wall properties, we applied Fourier-transformed infrared (FTIR) microspectroscopy to cells located in the middle part of the hypocotyl. The relative absorbance intensities for wavelengths related to the cell wall (from 830 to 1800 cm^-1^) were selected to assess the composition and status of various cell wall polysaccharides. FTIR analysis revealed significantly different patterns of relative absorption induced by low R:FR ratio after three days of treatment (**Fig. 2B and Fig. S1**). The main differences were observed between 1530 and 1200 cm^-1^ and between 1180 and 1030 cm^-1^ (**Fig. 2B)**. A relative decrease (at 1740 and between 1530-1200 cm^-1^) indicates lower levels of pectins and xyloglucans, as well as cellulose and lignin. On the other hand, an increase between 1180 and 1030 cm^-1^ indicates enrichement of pectins, xyloglucans, cellulose and arabinogalactan that can relate both to AGPs and rhamnogalacturonan I-type pectins (**Fig. 2B**). These results indicate major cell wall remodeling in response to low R:FR ratio. In order to obtain further insight into the changed cell wall components, we performed hierarchical clustering for the wavelengths that have previously been assigned to certain cell wall components (Kakuráková et al., 2000; Wilson et al., 2000; Mouille et al., 2003; Alonso-simón et al., 2011; Szymanska-Chargot and Zdunek, 2013; Largo-Gosens et al., 2014) (**Fig. 2C**). This revealed an overall enrichment for pectins with a low degree of methylesterification (de-esterified pectins), in addition to other pectins, arabinogalactan proteins, cellulose, and xyloglucans in response to low R:FR, while other wavelengths assigned to pectins, cellulose and xyloglucans showed opposite trends. Depending on the chemical bonds revealed by the wavelengths, different structural part of the polysaccharides are assessed such as backbone and side chains. This could explain opposite trends for wavelengths related to the same polysaccharide, which can be more abundant with reduced side chains for instance. Taken together, these results revealed major changes in cell wall biosynthesis and remodeling in response to low R:FR.

### Low R:FR impinges on expression of cell wall-related genes

In order to investigate transcriptional regulation of cell wall properties under low R:FR, we considered 40 gene families that are thought to be involved in synthesis and/or remodeling of cell wall components. The expression of a total of 824 genes was analyzed using an RNA sequencing dataset from a time course experiment under low R:FR conditions (Kohnen et al., 2016). In this set of genes, 544 were found to be expressed in seedlings grown under control conditions, among which 224 were regulated by low R:FR in hypocotyls (**Fig. 3A**). Only few genes were regulated in cotyledons or in both organs (26 and 16 genes, respectively) (**Fig. 3A**). Most genes were regulated at later stages (90 or 180 min after onset of light stimulus), and they were mostly up-regulated, in particular in hypocotyls (**Fig. 3A**). Modulated genes comprised functions related to (hemi)cellulose (network 1), pectin (network 2), and structural proteins (network 3) (**Fig. 3B**). The few genes that were induced at the early time points (15 or 45 min. after onset of the light stimulus) in hypocotyls are involved in cell wall remodelling (At1g49490:EXT, At5g47500:PME, At1g62760:PMEI, At5g02260:EXP, At5g57560:XTH, At1g02405:EXT, At3g10710:PME, **Tables S1A and S1B**). Considering the global influence of FR light on cell biosynthetic genes, the highest percentage related to pectins (network 2) with 96% of the genes expressed in seedlings, and 52% affected by low R:FR conditions. Taken together, these results indicate that cell wall remodelling is initiated within the first 15 minutes of low R:FR treatment, followed by general cell wall modifications involving both synthesis and remodeling of all cell wall constituents, in particular of pectin.

**Figure 3.**
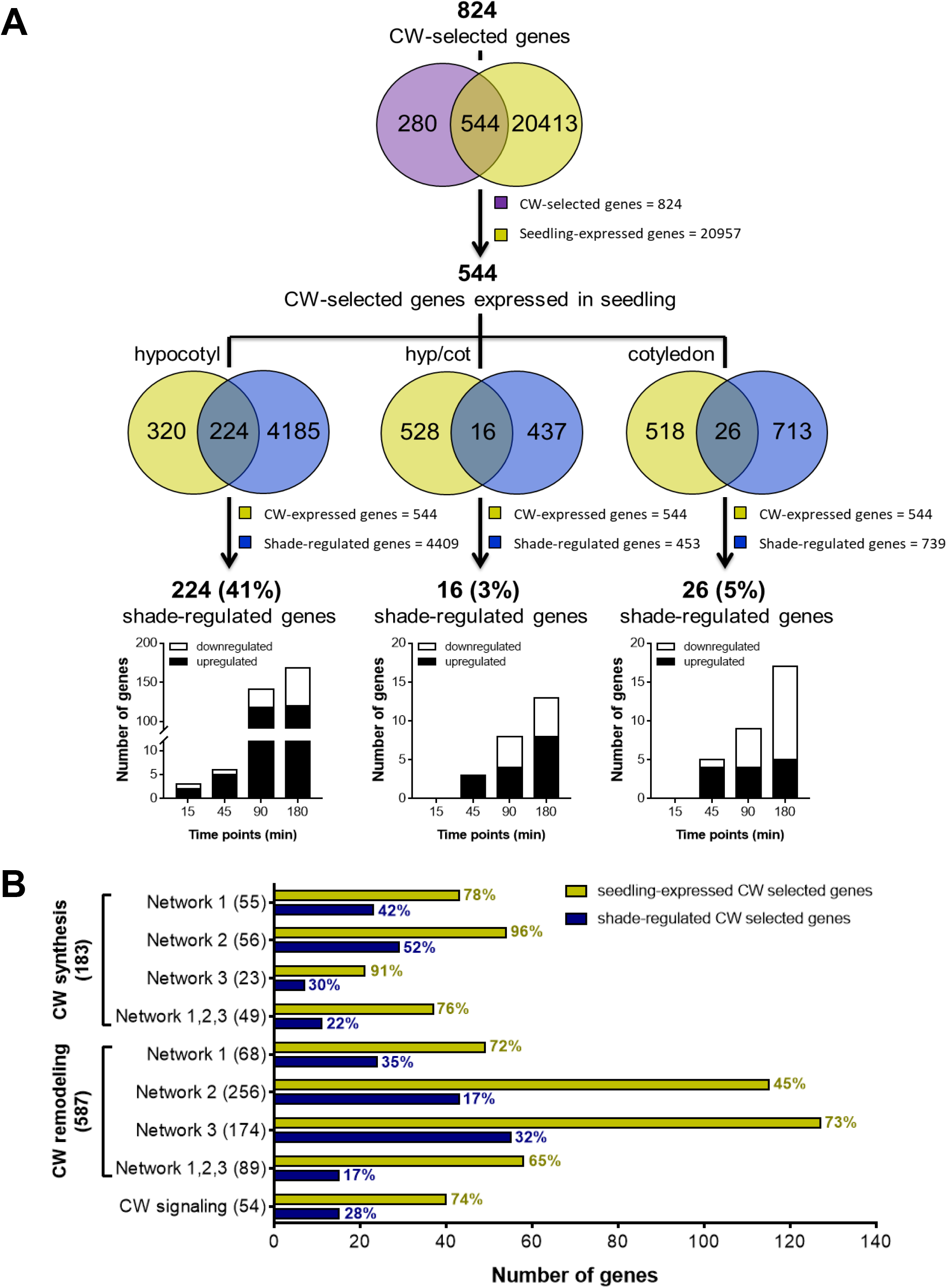
Low R:FR triggers changes in the expression of cell wall-related genes. **(A)** Number of cell wall-related genes identified as expressed in seedling and regulated by low R:FR in hypocotyl, cotyledon or both. From cell wall-selected genes and RNA sequencing data (Kohnen *et al*. 2016), Venn diagrams highlight cell wall-related genes expressed in seedling and that are regulated by low R:FR in hypocotyl, cotyledon or both. For each, number of up and down-regulated genes are shown along the kinetic of low R:FR treatment. Percentages of the low R:FR-regulated genes were determined according to the total of cell wall-related genes expressed in seedling. **(B)** Number of cell wall-related genes expressed in seedling and regulated by low R:FR classified according to their putative function in cell wall synthesis, remodeling and signaling as well as their related networks for synthesis and remodeling. Percentages were determined according to the total of cell wall-related genes classified for each condition (values between brackets). Network 1: cellulose and hemicelluloses ; Network 2: pectins ; Network 3: structural proteins ; CW: Cell Wall.

SAS is controlled by PIFs, hence, we interrogated previously published ChIP sequencing data (Kohnen et al., 2016) for interactions of PIF4 and PIF5 with the 224 genes regulated in the hypocotyl under low R:FR conditions. Indeed, 59 genes showed a direct interaction with PIF4 and/or PIF5 (**Fig. S2 and Table S1C**), including genes related to all three polymer networks, as well as to all processes including cell wall synthesis, remodelling, and signalling.

### Cell wall-related mutants reveal the role of cell wall components in low R:FR-induced growth

In order to gain insight into the mechanisms involved in low R:FR-induced growth, loss-of-function mutants defective in the three networks (cellulose, pectin, glycoproteins) were investigated for growth phenotypes under low R:FR treatment. We selected *xxt* mutants impaired in xyloglucan biosynthesis (network 1), mutants affected in pectin biosynthesis (*gaut, gatl* and *cgr*) and pectin remodeling (*pme* and *pmei*; network 2), and mutants affected in AGP biosynthesis (*galt, agp10c*, and *fla9*; network 3). All of these mutants were subjected to low R:FR treatment for three days and their hypocotyl growth response was normalized to the wild type. None of the single mutants in network 1 had a growth phenotype (**Fig. 4A**), possibly because of genetic redundancy, but the double mutants *xxt1xxt2* and *xxt2xxt5* showed significantly reduced hypocotyl elongation in response to low R:FR. In network 2 mutants, only *cgr2*, had a growth defect, which was exacerbated in the *cgr2cgr3* double mutant. Unexpectedely, mutants affected in network 3 generally grew longer than the wild type (**Fig. 4A**). Taken together, these results highlight the importance of xyloglucan and pectin in SAS, while some proteinaceous component of the cell walls appear to restricts hypocotyl elongation in the wild type.

**Figure 4.**
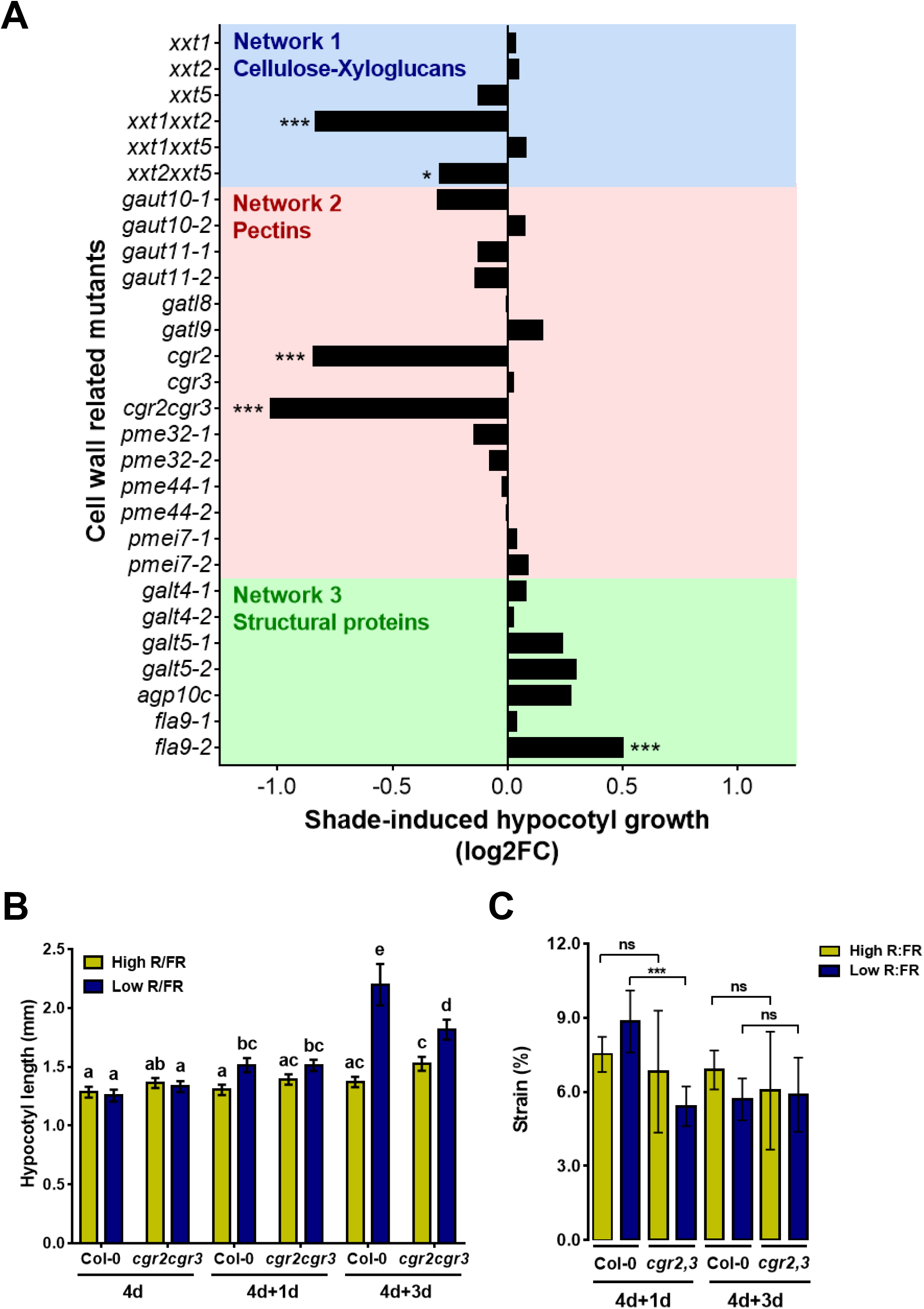
Analysis of mutants impaired in cell wall metabolism reveals importance of CGR2 and CGR3 in the regulation of the hypocotyl growth and the elastic properties under low R:FR. **(A)** Length of the hypocotyls in response to low R:FR treatment. Seedlings were grown for 4 days in high R:FR and then for 3 additional days in low R:FR. Data of growth induced by low R:FR during the 3 days were normalized against the wild-type and expressed in log2FC. Significant differences (p<0.05*, p<0.001***) were determined according to Student’s t-test. **(B)** Length of the hypocotyls in response to high and low R:FR treatments in Col-0 and *cgr2cgr3*. Seedlings were grown for 4 days in high R:FR before being transferred under low R:FR (blue bars) or kept in high R:FR (yellow bars) for 3 additional days. The bars show means in mm ± confidence intervals measured at 4 (4d), 5 (4d+1d) and 7 (4d+3d) days. Significant differences (indicated with letters) were determined according to one-way Anova followed by a multiple comparisons with Tukey’s test. **(C)** Elastic properties of hypocotyl assessed under high and low R:FR treatment for Col-0 and *cgr2cgr3*. Hypocotyls were frozen and thawed then subjected to cyclic loading at 5 mN of force and the strain compared to the data obtained for the wild-type seedlings in Figure 2. The average magnitude of strain incurred by seedlings grown in a high R:FR or low R:FR light regime after one (4d+1d) and three days (4d+3d) is shown. The bars show means in % ± SD (n>10 independent seedlings for Col-0 and n>5 independent seedlings for *cgr2cgr3*, at least five oscillations were made per seedling). Pairwise comparisons were made between the mutant and the wild-type using Welch t-test brackets indicated statistical tests that were made with significance p<0.1*, p<0.05** and p<0.01***.

Based on the induction of several homogalacturonan methyltransferases (*HGMT*), including the previously characterized HGMT genes *CGR2* and *CGR3* (Kim et al., 2015) (**Fig. S3**), and on the strong growth phenotype of the *cgr2cgr3* double mutant, (**Fig. 4A**), we investigated cell wall constituents in *cgr2cgr3* by FTIR analyses using wavelengths assigned to methylesterified (1740 cm^-1^) and demethylesterified (1630 cm^-1^) pectins (**Fig. S4**). Using these wavelengths we estimated the degree of methylesterification (DM), which was decreased by approximately 40% in *cgr2cgr3* compared to the wild type. In a time course experiment, *cgr2cgr3* was not affected in hypocotyl growth under control conditions (high R:FR), however, under low R:FR conditions, growth was reduced (**Fig. 4B**). Notably, *cgr2cgr3* reacted slower (4d+1d) and weaker (4d+3d) than the wild type (**Fig. 4B**). The growth defect of *cgr2cgr3* was particularly pronounced in the middle part of the hypocotyl that normally shows the strongest growth response (**Fig. S5**). This indicates that *cgr2cgr3* mutants have a defect in low R:FR-induced epidermal cell elongation.

We next assessed the mechanical properties of *cgr2cgr3* hypocotyls in response to low R:FR (**Fig. 4C**). Wild type hypocotyls had shown an increase in cell wall strain after 1d, and a decrease after 3d of FR treatment (**Fig. 2A**). In contrast, *cgr2cgr3* did not show a change in strain at either time point, indicating that cell wall remodeling is defective in the double mutant. To further address this aspect, we investigated cell wall composition of *cgr2cgr3* by FTIR microspectroscopy as in the wild type (**Fig. 2B**,**C**), in the range of wavelengths from 830 to 1800 cm^-1^ to obtain a cell wall fingerprint. This analysis was performed in the middle part of hypocotyls which shows the strongest growth increment under low R:FR (**Fig. 1B, Fig. S6**). Relative absorbances for Col-0 and *cgr2cgr3* did not show significant changes in response to low R:FR after the first day (**Fig. 5A; Figs. S6A**,**B upper panels**). However, 3 days after transfer, both wild type and *cgr2cgr3* showed significant changes of relative absorbances in response to low R:FR conditions **(Fig. 5A; Figs. S6A**,**B, lower panels**). Overall, the pattern of the significantly affected wavelengths were similar for both genotypes, but the differences were more pronounced in the wild type than *cgr2cgr3* (**Fig. 5B**).

**Figure 5.**
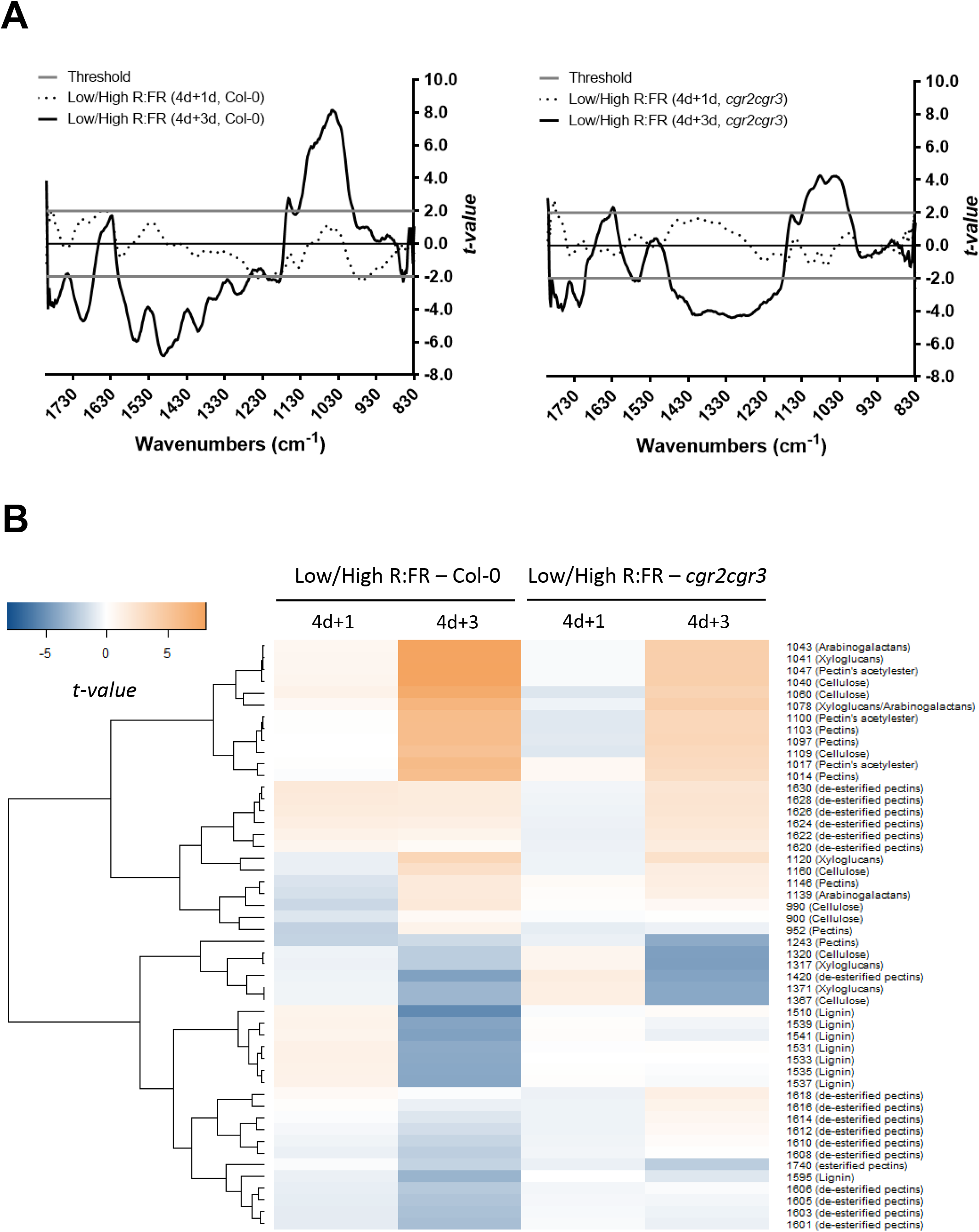
Changes of cell wall properties that occur in response to low R:FR are reduced in *cgr2cgr3*. **(A**,**B)** Cell wall properties in the middle part of the hypocotyl under high and low R:FR treatments for Col-0 and *cgr2cgr3*. Cell wall chemical bounds were analyzed by Fourrier-Transformed InfraRed (FTIR) microspectroscopy. For each hypocotyl, 6 spectra were collected in the middle part, avoiding the central cylinder, for at least 5 independent hypocotyls per condition. Baseline correction and data normalization were made for the absorbances between 1810 and 830 cm^-1^ (corresponding to the cell wall fingerprint, **see Supplemental Figure S6**). Pairwise comparison between high and low R:FR was made after 1 and 3 days treatments for Col-0 and *cgr2cgr3* and significant differences were identified using Student’s t-test for each wavelength. **(A)** All Student’s t-values were plotted against wavelengths with horizontal lines referring to significant threshold for p<0.05 for Col-0 (left panel) and *cgr2cgr3* (right panel). Student’s t-values above +2 or below -2 indicate respectively an enrichment or an impoverishment of cell wall components in low compared to high R:FR. **(B)** Student’s t-values for wavelengths assigned to cell wall components were used to build the heatmap with negative and positive t-values respectively represented by a range of colors from blue to orange.

The main differences beween the genotypes were observed in the range of wavelengths from 1630 to 1500 cm^-1^ with a decrease in the wild type but not in *cgr2cgr3* (**Fig. 5B**), and between 1180 to 1030 cm^-1^, where the wild type showed a stronger increase than the double mutant. These results suggest that a change in pectin methylesterification in *cgr2cgr3* results in secondary changes in cell wall composition (**Fig. 5B**).

## Discussion

While growth is simple to quantify, in particular in an organ like the hypocotyl that grows essentially in only one dimension, the underlying molecular mechanisms are extremely complex because they involve multiple regulatory levels such as hormonal and metabolic regulation, transcriptional activation of growth-related genes, and ultimately remodeling of the complex three-dimensional cell wall polymer network. Thanks to the ease of mutational analysis of growth, many of the upstream regulatory components in SAS have been identified (Procko et al., 2014; Ballaré and Pierik, 2017; Fiorucci and Fankhauser, 2017). A reduced R:FR indicative of neighbour proximity is primarily perceived by phyB in cotyledons and young leaves (Ballaré and Pierik, 2017; Fiorucci and Fankhauser, 2017). This leads to PIF-mediated induction of auxin production, which following transport to the hypocotyl promotes elongation (Procko et al., 2014; Ballaré and Pierik, 2017; Fiorucci and Fankhauser, 2017). PIFs can also directly regulate expression of hypocotyl specific genes as shown for genes encoding enzymes involved in plasma-membrane biogenesis (Ince et al., 2022). Our data suggests low R:FR induces extensive hypocotyl-specific regulation of genes encoding enzymes involved in cell wall biosynthesis and remodeling (**Fig. 3**), that ultimately results in in alteration of cell wall properties and induction of growth. However, due to the interdependency of cell wall components, it has been difficult to disentangle the role of individual cell wall components in growth. Here, we identified and characterized pectin methylesterification status as a central element in the growth phenomenon of Arabidopsis seedlings in the shade avoidance response.

Pectin consists mainly of homogalacturonan (HG) which represents a linear polymer of galacturonic acid (GalA) synthesized by galacturonosyltransferases (GAUTs) and GAUT-like (GATL) enzymes in the Golgi apparatus (Atmodjo et al., 2013). HG is subsequently methylesterified by HGMTs before secretion into the cell wall, where it can be selectively demethylesterified by pectin methylesterases (PMEs) (Sénéchal et al., 2014b). Thus, PME activity adjusts the degree of methylesterification (DM), which in turn modulates cell wall mechanical properties (Peaucelle et al., 2011; Wang et al., 2020). Furthermore, PME-mediated demethylesterification can expose HG to pectin-degrading enzymes such as polygalacturonases (PGs) and pectate lyases-like (PLLs) (Sénéchal et al., 2014b). Pectin degradation by these enzymes can contribute to cell wall loosening. There is evidence for a functional role of pectin metabolism in the regulation of growth (Bouton, 2002; Mouille et al., 2007) (Pelletier et al., 2010; Guénin et al., 2011; Wolf et al., 2012b; Sénéchal et al., 2014a) (Wang et al., 2010; Xiao et al., 2014; Rui et al., 2017), however, the high degree of redundancy in cell-wall remodeling enzymes has complicated the analysis. For example, there are 66 and 76 genes, respectively, that encode PMEs and PME inhibitors (PMEIs), allowing for extensive compensatory responses upon genetic or pharmacological interference.

A mechanistic understanding of growth phenomena requires the combined use of genetic, analytic, and biomechanical methods to identify the causal elements in cell growth. Using FTIR analysis, we document changes in cell wall composition in response to changes in the R:FR ratio, which correlate with accelerated growth. We identified two homogalacturonan-methyltransferase (HGMT) genes (*CGR2* and *CGR3*) that are required for neighbour proximity-induced growth. Biomechanical assays showed that *CGR2* and *CGR3* contribute to cell wall extensibility at the onset of growth. Previous studies used atomic force microscopy to analyze cell walls, which meausures properties perpendicular to the direction of growth, to detect differences in mechanical properties in plants with modified pectin that correlated with growth (Peaucelle et al., 2008; Peaucelle et al., 2011; Braybrook and Peaucelle, 2013; Peaucelle et al., 2015). In this study we were able to measure differences in mechanical properties using an extensometer which measures properties in the direction of growth. Both approaches show a correlation between modifying pectin chemistry, changes in cell wall mechanical properties and growth, supporting a role of pectin in growth regulation. Further work is required to understand the nature of this regulation and how it relates to the other cell wall components (Coen and Cosgrove, 2023).

## Materials and Methods

Information on the biological material and methods is available in the supplemental data **Acknowledgement**

## Acknowledgement

The authors wish to gratefully thank Prof. Kenneth Keegstra, Prof. Debra Mohnen, Prof. Frederica Brandizzi and Prof. Jérôme Pelloux respectively for sharing seeds of the *xxt, gaut, cgr* and *pme/pmei* mutants used in this study. We also thank Dr. Christiane Nawrath, Dr. Sylvester Mazurek and Dr. Gregory Mouille for their help in the analysis of the FTIR microspectroscopy. We thank the Swiss initiative in systems biology (SystemX.ch) and the University of Lausanne for the funding support. We nicely thank Prof. Cris Kuhlemeier for contributing to the funding acquisition and the co-supervising of the “Plant Growth 2 in a Changing Environment” project funded by the Swiss initiative in systems biology (SystemX.ch). We also thank him for his help in the project handing and for hosting Sarah Robinson, postdoc in his lab during the project.

## Literature cited

Alonso-simón A, García-angulo P, Mélida H, Encina A, Álvarez JM, Acebes JL (2011) The use of FTIR spectroscopy to monitor modifications in plant cell wall architecture caused by cellulose biosynthesis inhibitors The use of FTIR spectroscopy to monitor modifications in plant cell wall architecture caused by cellulose biosynthesis inhibi. Plant Signal Behav 6: 1104–1110

Atmodjo MA, Hao Z, Mohnen D (2013) Evolving Views of Pectin Biosynthesis. Annu Rev Plant Biol 64: 747–779

Ballaré CL, Pierik R (2017) The shade-avoidance syndrome: Multiple signals and ecological consequences. Plant Cell Environ 40: 2530–2543

Bashline L, Lei L, Li S, Gu Y (2014) Cell wall, cytoskeleton, and cell expansion in higher plants. Mol Plant 7: 586–600

Bouton S (2002) QUASIMODO1 Encodes a Putative Membrane-Bound Glycosyltransferase Required for Normal Pectin Synthesis and Cell Adhesion in Arabidopsis. Plant Cell Online 14: 2577–2590

Braybrook SA, Peaucelle A (2013) Mechano-Chemical Aspects of Organ Formation in Arabidopsis thaliana: The Relationship between Auxin and Pectin. PLoS One. doi: 10.1371/journal.pone.0057813

Coen E, Cosgrove DJ (2023) The mechanics of plant morphogenesis. Science 379: eade8055

Cosgrove DJ (2005) Growth of the plant cell wall. Nat Rev Mol Cell Biol 6: 850–861

Fiorucci AS, Fankhauser C (2017) Plant Strategies for Enhancing Access to Sunlight. Curr Biol 27: R931–R940

Galvão VC, Fankhauser C (2015) Sensing the light environment in plants: Photoreceptors and early signaling steps. Curr Opin Neurobiol 34: 46–53

Gommers CMM, Visser EJW, Onge KRS, Voesenek LACJ, Pierik R (2013) Shade tolerance: When growing tall is not an option. Trends Plant Sci 18: 65–71

Guénin S, Mareck A, Rayon C, Lamour R, Assoumou Ndong Y, Domon JM, Sénéchal F, Fournet F, Jamet E, Canut H, et al (2011) Identification of pectin methylesterase 3 as a basic pectin methylesterase isoform involved in adventitious rooting in Arabidopsis thaliana. New Phytol 192: 114–126

Hijazi M, Velasquez SM, Jamet E, Estevez JM, Albenne C (2014) An update on posttranslational modifications of hydroxyproline-rich glycoproteins: toward a model highlighting their contribution to plant cell wall architecture. Front Plant Sci 5: 1–10

Ince YÇ, Krahmer J, Fiorucci AS, Trevisan M, Galvão VC, Wigger L, Pradervand S, Fouillen L, Van Delft P, Genva M, et al (2022) A combination of plasma membrane sterol biosynthesis and autophagy is required for shade-induced hypocotyl elongation. Nat Commun. doi: 10.1038/s41467-022-33384-9

Kakuráková M, Capek P, Sasinkova V, Wellner N, Ebringerova A, Kac M (2000) FT-IR study of plant cell wall model compounds : pectic polysaccharides and hemicelluloses. Carbohydr Polym 43: 195–203

Kim S-J, Held MA, Zemelis S, Wilkerson C, Brandizzi F (2015) CGR2 and CGR3 have critical overlapping roles in pectin methylesterification and plant growth in Arabidopsis thaliana. Plant J 82: 208–20

Kohnen M V., Schmid-Siegert E, Trevisan M, Petrolati LA, Sénéchal F, Müller-Moulé P, Maloof J, Xenarios I, Fankhauser C (2016) Neighbor detection induces organspecific transcriptomes, revealing patterns underlying hypocotyl-specific growth. Plant Cell 28: 2889–2904

Largo-Gosens A, HernÃ¡ndez-Altamirano M, GarcÃ-a-Calvo L, Alonso-SimÃn A, Ãlvarez J, Acebes JL (2014) Fourier transform mid infrared spectroscopy applications for monitoring the structural plasticity of plant cell walls. Front Plant Sci 5: 1–15

Legris M, Ince YÇ, Fankhauser C (2019) Molecular mechanisms underlying phytochrome-controlled morphogenesis in plants. Nat Commun 10: 1–15

Mouille G, Ralet MC, Cavelier C, Eland C, Effroy D, Hématy K, McCartney L, Truong HN, Gaudon V, Thibault JF, et al (2007) Homogalacturonan synthesis in Arabidopsis thaliana requires a Golgi-localized protein with a putative methyltransferase domain. Plant J 50: 605–614

Mouille G, Robin S, Lecomte M, Pagant S, Höfte H (2003) Classification and identification of Arabidopsis cell wall mutants using Fourier-Transform InfraRed (FT-IR) microspectroscopy. Plant J 35: 393–404

Nguema-Ona E, VicrÃ©-Gibouin M, GottÃ© M, Plancot B, Lerouge P, Bardor M, Driouich A (2014) Cell wall O-glycoproteins and N-glycoproteins: aspects of biosynthesis and function. Front Plant Sci 5: 1–12

Pauly M, Keegstra K (2016) Biosynthesis of the Plant Cell Wall Matrix Polysaccharide Xyloglucan. Annu Rev Plant Biol 67: 235–259

Peaucelle A, Braybrook SA, Le Guillou L, Bron E, Kuhlemeier C, Höfte H (2011) Pectin-induced changes in cell wall mechanics underlie organ initiation in Arabidopsis. Curr Biol 21: 1720–1726

Peaucelle A, Louvet R, Johansen JN, Höfte H, Laufs P, Pelloux J, Mouille G (2008) Arabidopsis Phyllotaxis Is Controlled by the Methyl-Esterification Status of Cell-Wall Pectins. Curr Biol 18: 1943–1948

Peaucelle A, Wightman R, Höfte H (2015) The Control of Growth Symmetry Breaking in the Arabidopsis Hypocotyl. Curr Biol 25: 1746–1752

Pedmale U V., Huang SSC, Zander M, Cole BJ, Hetzel J, Ljung K, Reis PAB, Sridevi P, Nito K, Nery JR, et al (2016) Cryptochromes Interact Directly with PIFs to Control Plant Growth in Limiting Blue Light. Cell 164: 233–245

Pelletier S, Van Orden J, Wolf S, Vissenberg K, Delacourt J, Ndong YA, Pelloux J, Bischoff V, Urbain A, Mouille G, et al (2010) A role for pectin de-methylesterification in a developmentally regulated growth acceleration in dark-grown Arabidopsis hypocotyls. New Phytol 188: 726–739

Pierik R, De Wit M (2014) Shade avoidance: Phytochrome signalling and other aboveground neighbour detection cues. J Exp Bot 65: 2815–2824

Procko C, Crenshaw CM, Ljung K, Noel JP, Chory J (2014) Cotyledon-generated auxin is required for shade-induced hypocotyl growth in brassica rapa. Plant Physiol 165: 1285–1301

Pucciariello O, Legris M, Costigliolo C, José M, Esteban C, Dezar C, Vazquez M, Yanovsky MJ, Finlayson SA, Prat S, et al (2018) Rewiring of auxin signaling under persistent shade. Proc Natl Acad Sci 2–7

de Reuille PB, Routier-Kierzkowska AL, Kierzkowski D, Bassel GW, Schüpbach T, Tauriello G, Bajpai N, Strauss S, Weber A, Kiss A, et al (2015) MorphoGraphX: A platform for quantifying morphogenesis in 4D. Elife 4: 1–20

Robinson S, Huflejt M, Barbier de Reuille P, Braybrook SA, Schorderet M, Reinhardt D, Kuhlemeier C (2017) An automated confocal micro-extensometer enables in vivo quantification of mechanical properties with cellular resolution. Plant Cell tpc.00753.2017

Rui Y, Xiao C, Yi H, Kandemir B, Wang JZ, Puri VM, Anderson CT (2017) POLYGALACTURONASE INVOLVED IN EXPANSION3 Functions in Seedling Development, Rosette Growth, and Stomatal Dynamics in Arabidopsis thaliana. Plant Cell 29: 2413–2432

Sasidharan R, Chinnappa CC, Staal M, Elzenga JTM, Yokoyama R, Nishitani K, Voesenek Lacj, Pierik R (2010) Light Quality-Mediated Petiole Elongation in Arabidopsis during Shade Avoidance Involves Cell Wall Modification by Xyloglucan Endotransglucosylase/Hydrolases. Plant Physiol 154: 978–990

Sasidharan R, Chinnappa CC, Voesenek Lacj, Pierik R (2008) The Regulation of Cell Wall Extensibility during Shade Avoidance: A Study Using Two Contrasting Ecotypes of Stellaria longipes. Plant Physiol 148: 1557–1569

Sasidharan R, Pierik R (2010) Cell wall modification involving XTHs controls phytochrome-mediated petiole elongation in Arabidopsis thaliana. Plant Signal Behav 5: 1491–1492

Sénéchal F, Graff L, Surcouf O, Marcelo P, Rayon C, Bouton S, Mareck A, Mouille G, Stintzi A, Höfte H, et al (2014a) Arabidopsis PECTIN METHYLESTERASE17 is co-expressed with and processed by SBT3.5, a subtilisin-like serine protease. Ann Bot 114: 1161–1175

Sénéchal F, Wattier C, Rustérucci C, Pelloux J (2014b) Homogalacturonan-modifying enzymes: Structure, expression, and roles in plants. J Exp Bot 65: 5125–5160

Showalter AM, Basu D (2016) Extensin and Arabinogalactan-Protein Biosynthesis: Glycosyltransferases, Research Challenges, and Biosensors. Front Plant Sci 7: 1–9

Szymanska-Chargot M, Zdunek A (2013) Use of FT-IR Spectra and PCA to the Bulk Characterization of Cell Wall Residues of Fruits and Vegetables Along a Fraction Process. Food Biophys 8: 29–42

Wang H, Guo Y, Lv F, Zhu H, Wu S, Jiang Y, Li F, Zhou B, Guo W, Zhang T (2010) The essential role of GhPEL gene, encoding a pectate lyase, in cell wall loosening by depolymerization of the de-esterified pectin during fiber elongation in cotton. Plant Mol Biol 72: 397–406

Wang X, Wilson L, Cosgrove DJ (2020) Pectin methylesterase selectively softens the onion epidermal wall yet reduces acid-induced creep. J Exp Bot 71: 2629–2640

Wilson RH, Smith AC, Kac M, Saunders PK, Wellner N, Waldron KW (2000) The Mechanical Properties and Molecular Dynamics of Plant Cell Wall Polysaccharides Studied by Fourier-Transform Infrared Spectroscopy 1. Plant Physiol 124: 397–405

de Wit M, Galvão VC, Fankhauser C (2016) Light-Mediated Hormonal Regulation of Plant Growth and Development. Annu Rev Plant Biol 67: 513–537

de Wit M, Lorrain S, Fankhauser C (2014) Auxin-mediated plant architectural changes in response to shade and high temperature. Physiol Plant 151: 13–24

Wolf S, Hématy K, Höfte H (2012a) Growth Control and Cell Wall Signaling in Plants. Annu Rev Plant Biol 63: 381–407

Wolf S, Mravec J, Greiner S, Mouille G, Höfte H (2012b) Plant cell wall homeostasis is mediated by brassinosteroid feedback signaling. Curr Biol 22: 1732–1737

Xiao C, Somerville C, Anderson CT (2014) POLYGALACTURONASE INVOLVED IN EXPANSION1 Functions in Cell Elongation and Flower Development in Arabidopsis. Plant Cell 26: 1018–1035

